# Decoding the Human Brain during Intelligence Testing

**DOI:** 10.1101/2025.04.01.646660

**Authors:** Jonas A. Thiele, Joshua Faskowitz, Olaf Sporns, Adam Chuderski, Rex Jung, Kirsten Hilger

**Affiliations:** Department of Psychology I, Würzburg University, Marcusstr. 9-11, Würzburg D-97070, Germany; Department of Psychological and Brain Sciences, Indiana University, 1101 E. 10th St., Bloomington, IN 47405-7007, USA; Centre for Cognitive Science, Jagiellonian University, Ingardena 3, 30-060 Krakow, Poland; The University of New Mexico, Department of Psychology, Logan Hall MSC03-2220, 1 University of New Mexico, Albuquerque, NM 87131-0001, USA; Department of Psychology, Differential Psychology, Personality Psychology and Psychological Diagnostics, Vinzenz Pallotti University Vallendar, Germany

**Keywords:** Intelligence, Raven’s Progressive Matrices (RPM), EEG, fMRI, Functional Connectivity, Connector Hubs

## Abstract

Understanding the brain mechanisms underlying complex human cognition is a major objective in neuroscience. Previous studies have identified neural correlates of intelligence at different temporal and spatial scales using functional magnetic resonance imaging (fMRI) and electroencephalography (EEG) separately. This study treats intelligence as a multilayer phenomenon across temporal and spatial scales and examines how the connectedness of brain regions as well as the complexity of multilayer brain dynamics relates to performance in an established intelligence test. Graph-theoretical analyses of fMRI-derived functional connectivity (*N*=67) revealed that the connectedness of frontal and parietal regions was associated with individual performance. Further, multiscale entropy analyses of EEG signals (*N*=131) disclosed that higher test scores were linked to more complex long-range processes and, at a trend level, to less complex short-range processes. These findings support the Multilayer Processing Theory, proposing the interplay between flexible long-range and modular short-range processes as the neural bases of intelligence.

## Introduction

Intelligence captures a person’s individual level of general cognitive ability. It predicts important life outcomes, including academic achievement, health, and longevity^1,2^. For more than a hundred years, research has been seeking its biological bases, mostly focusing on the central nervous system^3,4^. While the scientific study of intelligence has contributed valuable insights into human cognition, it is important to acknowledge that intelligence testing has also been historically misused, particularly in the context of eugenics. Recognizing this legacy is essential for ensuring ethically grounded research in this domain.

Across the last decade, numerous associations between individual intelligence scores and characteristics of brain structure and function have been identified with techniques like functional magnetic-resonance imaging (fMRI) and electroencephalography (EEG)^5^. The Parieto-Frontal Integration Theory (P-FIT) is one of the most influential theories regarding the neural basis of intelligence. It proposes that both structural and functional characteristics of circumscribed brain regions, particularly the communication between frontal and parietal brain regions, is crucial to differences in intelligence^6,7^. More recent proposals expanded the P-FIT, including the Brain Connectivity Model (BCM)^8^ and the Network Neuroscience Theory of Intelligence (NNT)^9^. While the BCM presents an extensive summary of empirical studies but lacks mechanistic hypotheses, the NNT proposes the flexibility to transition between different brain network states as critical to explain variation in intelligence^9, 10^. However, empirical evidence supporting the NNT is limited^11^. The recently proposed Multilayer Processing Theory of Intelligence (MLPT)^12^ offers a conceptual framework that bridges the localizationist and dynamic perspectives by conceptualizing intelligence as emerging from hierarchical layers of processing: global long-range processes in integrative networks, including fronto-parietal brain regions, operating at coarser temporal scales, and short-range subprocesses within smaller neuronal assemblies, taking place on finer temporal scales. These latter more specialized short-range processes are coordinated by the former long-range network processes. Intelligence-related processes are proposed to manifest differently across temporal scales, with higher intelligence resulting from simpler short-range processes and more flexible, complex long-range processes. However, this theory has not yet been empirically tested, so far.

In general, coarse-scale global network flexibility might be based on functionally well-connected brain regions (connector hubs)^13^ orchestrating changes of activity patterns (network states) to meet external demands^9,14^. Previous fMRI research supports the critical role of network efficiency and flexibility for human behavior and thought^15^. In resting-state functional connectivity studies, variations in the connectedness of hub regions^16,17^ as well as the stability and flexibility of brain network organization over time were identified as significant correlates of individual differences in intelligence^18–21^. Further, studies assessing functional connectivity during task demands suggest that adapting brain networks fluidly^22^ and efficiently^23–25^ to different demands, supports intelligence-related cognitive processing. While fMRI connectivity studies provide insight into brain communication patterns at coarse temporal scales (e.g., via slow, correlated BOLD fluctuations), for testing assumptions of the MLPT, those need to be complemented with characteristics of faster scale brain dynamics, e.g., brain oscillations assessed via EEG.

One aspect of the MLPT is that higher intelligence emerges from the simplicity of local short-range processes and the flexibility of global long-range processes. Those characteristics might be reflected in neural signal complexity, which relates to information richness^26,27^ and dynamic flexibility^28^. As lower time scales are proposed to be linked to short-range and higher scales to long-range processes^29–32^, low complexity at shorter scales might indicate simpler processing, while higher complexity at longer scales might reflect greater flexibility. Indeed, less complexity in short-range processes^33,34^ and higher complexity in long-range processes^33^ has previously been linked to higher intelligence.

Importantly, most of these previous connectivity-based fMRI studies and complexity-based EEG studies were based on brain activity recorded during resting state or during tasks requiring little cognitive effort. Although tasks with increasing difficulty have been considered^35,36^, research on tasks directly linked to intelligence-related cognitive processing^37,38^ remains rare^25,39^. Studies during performing intelligence tests - such as the Raven Progressive Matrices (RPM) - might provide more insight.

Using fMRI recorded during RPM, early studies observed associations between individual differences in intelligence and variations in brain activity in frontal and (posterior) parietal cortex^40–42^, in the insula^40^ as well as in regions of the default mode network (DMN)^43^. EEG recorded during RPM demonstrated associations between intelligence and brain activity (power) in multiple frequency bands^44^, relations between intelligence scores and differences in delta-gamma^45^ and theta-gamma cross-frequency coupling^46^, and revealed connectivity strength (between the left precentral gyrus and left dorsal cingulate) as being related to intelligence^47^. However, these studies provide no information about the flexibility or complexity of involved multiscale processes and are thus limited with respect to the MLPT’s major claims: Intelligence relates to more flexible long-range processes at coarser time scales, coordinating simpler short-range processes at finer time scales within distinctive regions.

In this study, fMRI and EEG analysis are combined to investigate characteristics of long-range and short-range processes in their association with intelligence, thereby addressing key assumptions of the MLPT. Specifically, we obtained: a) spatial information about long-range processes with graph-theoretical centrality measures applied to fMRI data, and b) information about the complexity (or simplicity) of long- and short-range processes with multiscale entropy (MSE) analyses of EEG data measured during an established intelligence test.

## Results

### Intelligence

To investigate intelligence-related neural processes across temporal and spatial scales, we analyzed datasets from two laboratories. The first (Sample 1) includes fMRI data from 67 participants and was used to examine slow functional communication and regional involvement with graph-theoretical centrality measures, i.e., network degree and participation coefficient. The second (Sample 2) comprises EEG data from 131 participants and was used to capture higher-temporal dynamics via MSE analysis. Both datasets include recordings from resting-state and RPM performance. RPM-specific cognitive processing was analyzed after subtraction of intrinsic (resting-state) centrality or entropy measures from the RPM measures (see Methods for details). In both samples, RPM sum scores were approximately normally distributed (*Supplement* Fig. S1). In Sample 1, scores ranged from 9.6 to 30 with a mean of 21.65, a standard deviation of 4.48 and they were not significantly correlated with age (*rho* = 0.16, *p* = 0.197), sex (*rho* = −0.19, *p* = 0.114) or framewise displacement (*rho* = −0.12, *p* = 0.354). In Sample 2, RPM scores ranged between 3 and 31 with a mean of 20.41 and a standard deviation of 5.21. The scores were significantly negatively correlated with age (*rho* = −0.24, *p* = 0.005), while no significant correlation with sex (*rho* = 0.07, *p* = 0.453) or the number of removed epochs (*rho* = −0.15, *p* = 0.092) was observed. Note that, as only 30 items had to be solved in Sample 1, whereas the test in Sample 2 consisted of 36 items, we multiplied the original scores of Sample 1 by a factor of 36/30 to allow for better comparability. The scores listed above, as well as those illustrated in Supplementary Figure S1, refer to the values after this adjustment.

### Functional MRI-based brain network centrality

Group-average functional connectivity matrices of the resting state, the RPM test as well as the difference between RPM and resting-state functional connectivity are shown in *Supplement* Figs S2-S4. Replicating previous studies that compared resting-state functional connectivity with functional connectivity obtained during other tasks, task performance led to a decrease in connectivity in primary sensory unimodal areas, while increased connectivity was observed in multimodal associative regions, as compared to resting state ^24^. Descriptively, this increase was strongest for connectivity between the default mode network and the ventral attention network.

Group-average brain maps of both graph-theoretical centrality measures, i.e., degree and participation coefficient, obtained during resting state and during RPM as well as their rest-task difference are displayed in Figures 1 and 2. During RPM performance, degree was descriptively higher in bilateral regions of the medial and dorsolateral prefrontal cortex, in wide-spread regions of the medial parietal cortex as well as in bilateral areas around the temporo-parietal junction areas (TPJ), while it was slightly lower in some visual, temporal and motor regions compared to resting state. Regarding the participation coefficient, a similar but more localized pattern was observed when transitioning from resting state to RPM with increases in the dorsolateral, dorsomedial, and ventromedial prefrontal cortices, and in bilateral TPJs including parts of the inferior partial lobules (IPL) and the superior temporal sulci (STS). Spatial correlations between both graph-theoretical centrality measures are presented in the Supplement.

**Figure 1.**
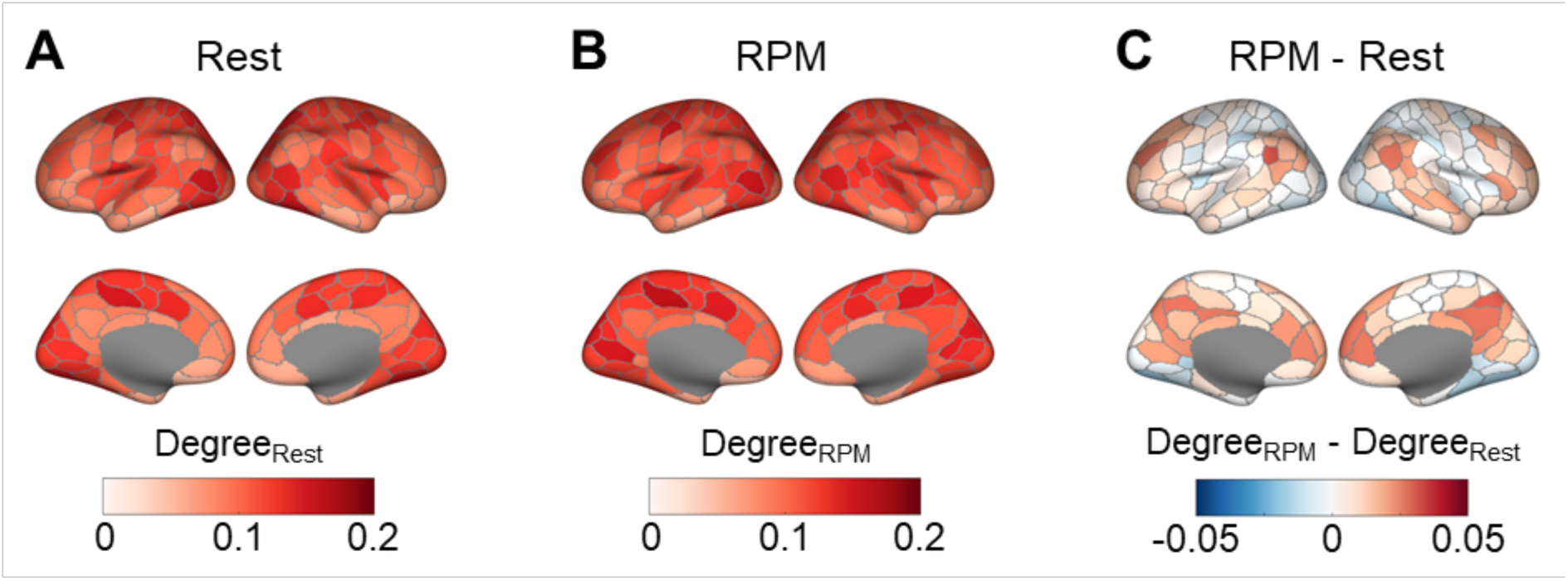
Degree derived from fMRI and averaged across all 67 participants of Sample 1. A) Degree during resting state, B) degree during RPM performance, C) difference in degree between RPM performance and resting state.

**Figure 2.**
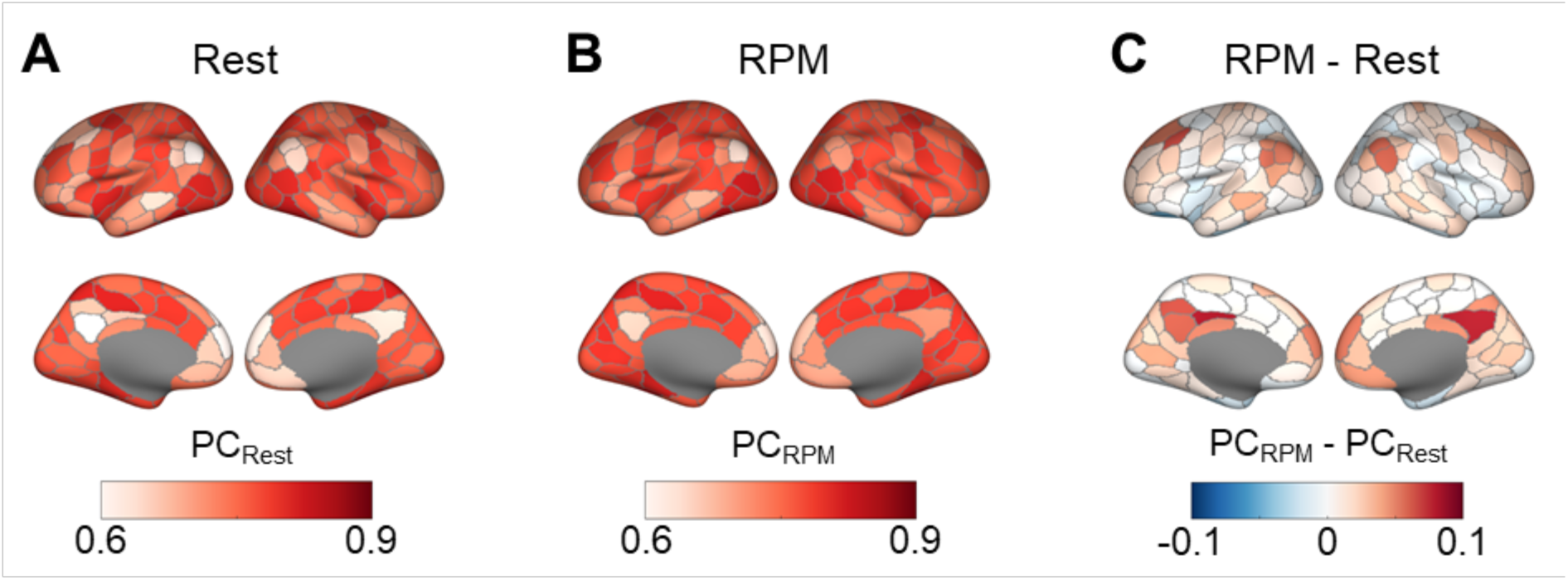
Participation coefficient (PC) derived from fMRI and averaged across all 67 participants of Sample 1. A) PC during resting state, B) PC during RPM performance, C) difference in PC between RPM performance and resting state.

### Electroencephalography-based multiscale entropy

Aggregated across all electrodes, group-average MSE obtained during resting state and during RPM are displayed in Figure 3. Global MSE was significantly higher during RPM than during resting state for all time scales above 2. This difference increased at higher temporal scales, suggesting an increase in the complexity of long-range processes.

**Figure 3.**
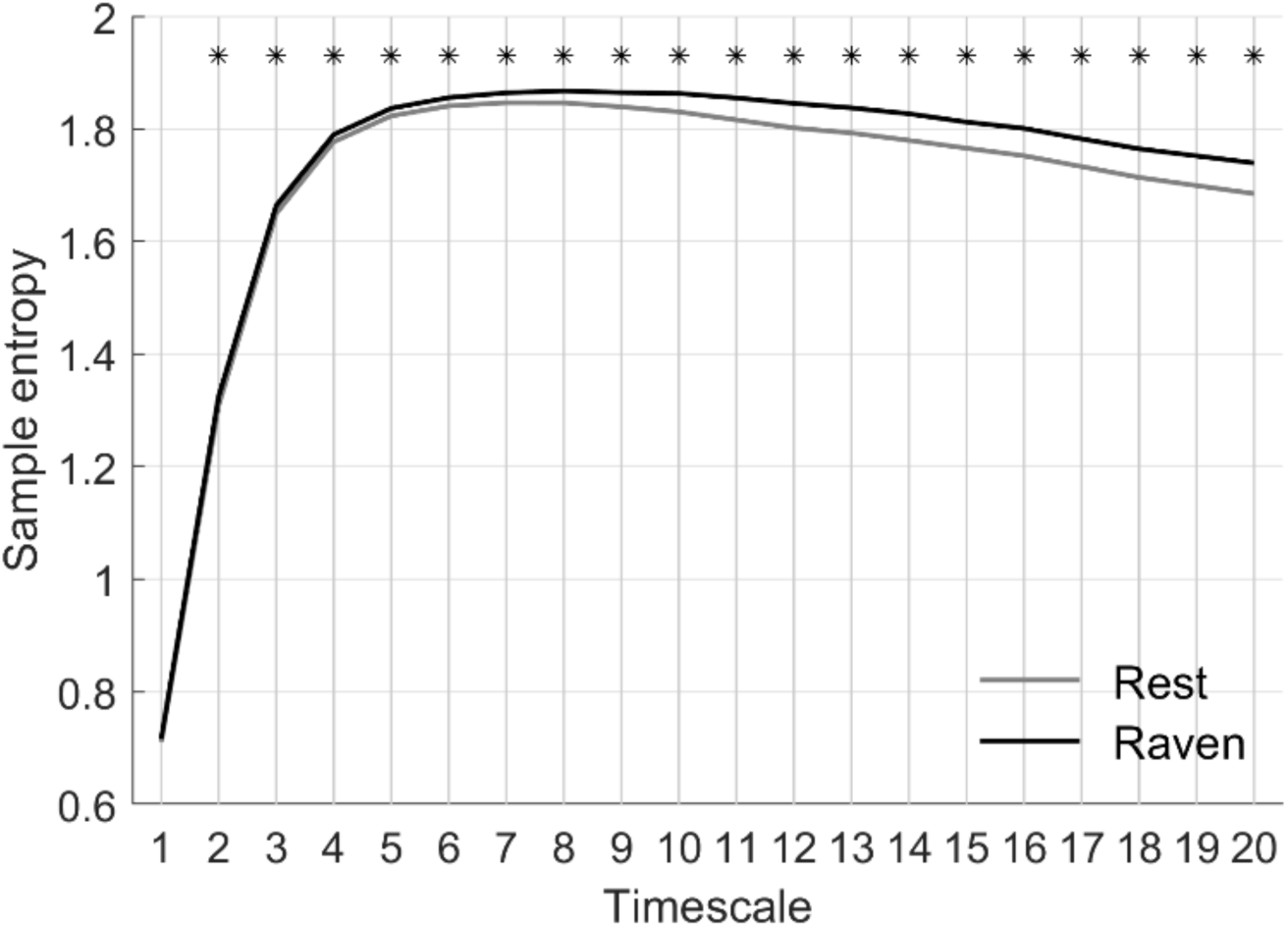
Multiscale entropy (MSE) derived from EEG and averaged across the 131 participants of Sample 2 and all 64 electrodes. The black line represents the group mean MSE during RPM performance, while the gray line indicates the group mean MSE during rest, averaged across participants and electrodes. Statistical significance at each timescale was assessed using paired *t*-tests across subjects, with *p*-values corrected for multiple comparisons using the False Discovery Rate (FDR) procedure. Small black asterisks above the curves indicate timescales where the difference between resting-state and RPM performance was significant after FDR correction (*p* < 0.05).

Group-average MSE for single electrodes are displayed in Figures 4 and reveal that the increase in signal complexity is primarily driven by temporal, frontal and parietal channels. While the increased entropy in T7 and T8 might reflect enhanced motion due to the execution of RPM-related button presses, higher entropy in frontal and parietal electrodes might reflect an increasing complexity of long-range processing of underlying brain regions.

**Figure 4.**
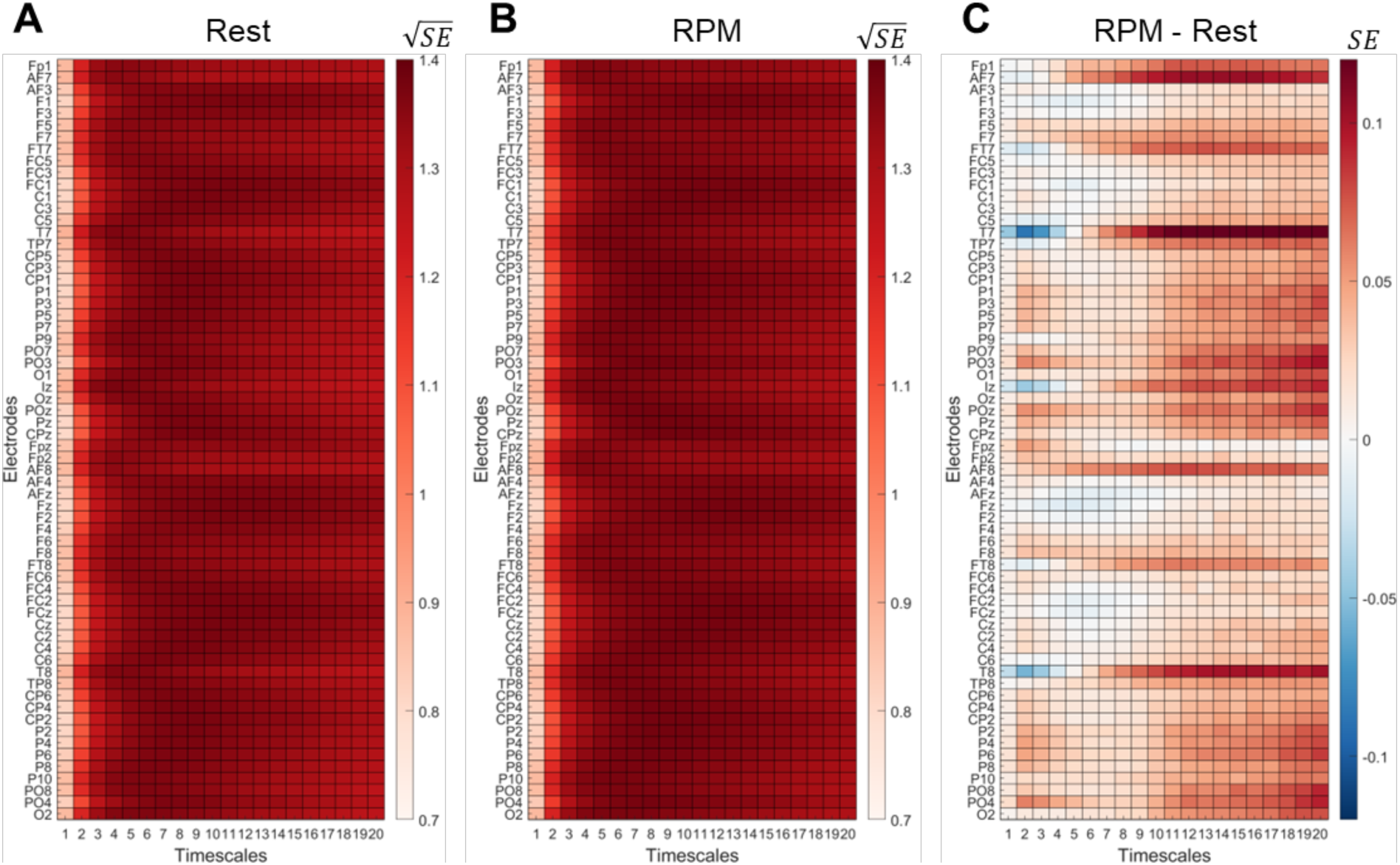
Electrode-specific multiscale entropy (MSE) derived from EEG and averaged across the 131 participants of Sample 2. A) MSE during resting state, B) MSE during RPM performance, C) difference in MSE between RPM performance and resting state. A) and B) show square-root-transformed MSE values for the rest and RPM conditions, respectively, to enhance visual differentiation of values across timescales. C) shows the raw (non-transformed) difference in MSE between RPM and resting state. Warmer colors in the difference plot indicate higher MSE during the RPM compared to rest, while cooler colors indicate the opposite. Each row corresponds to one EEG electrode, and each column to a timescale (1–20). SE, Sample entropy.

### Intelligence is associated with fMRI-based connectivity between frontal and parietal brain modules

In the fMRI sample, degree demonstrated a descriptive trend of mainly positive associations with individual intelligence scores in bilateral somatomotor, left lateral prefrontal and parietal regions as well as in regions of the medial prefrontal cortex (Figure 5, left panel). This trend was strongest in the left dorsolateral prefrontal cortex, in a region of the left ventro-lateral prefrontal cortex, in two regions of the left IPL (one of them belonging to the left TPJ) and in two bilateral somatomotor clusters. However, none of these relationships reached significance when controlling for the number of network nodes for which measures were calculated (200 cortical brain regions; 0.25 ≤ *rho* ≤ 0.31, all *p* > 0.05, controlled for age, sex, and head motion). Similar results were observed for varying functional connectivity thresholds (*Supplement* Figs. S5 and S6).

**Figure 5.**
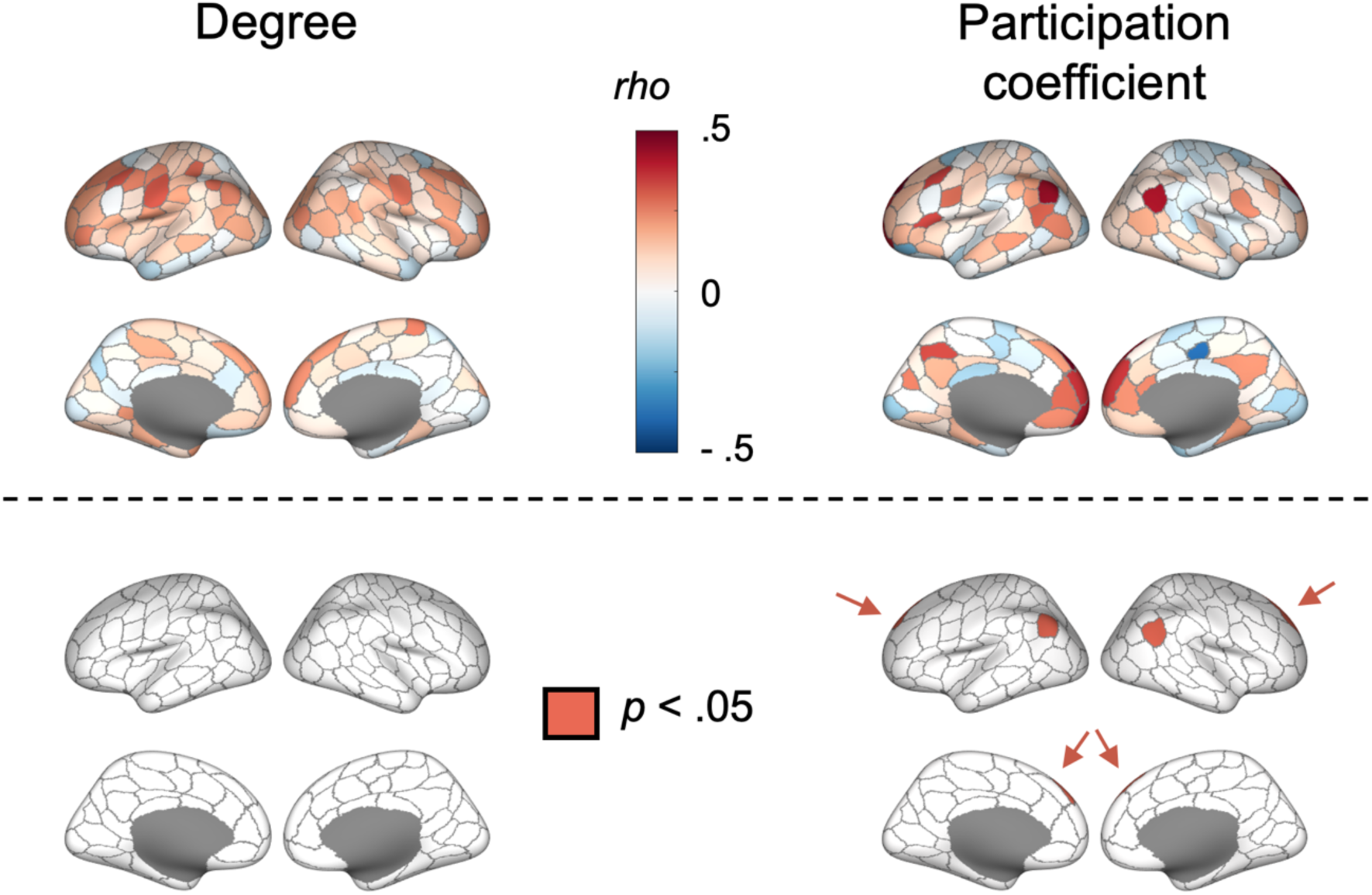
Intelligence is associated with fMRI-based centrality measures during intelligence testing. Associations between individual intelligence scores and variations in degree (left) and participation coefficient (right) in 200 cortical brain regions as assessed with fMRI during performing the RPM. Upper panels show the Spearman correlation rho controlled for age, sex, and head motion. In lower panels, areas with *p*-values < .05, indicating significance (corrected for the number of network nodes with FDR, i.e., 200 brain regions) are colored in red (irrespective of whether associations with intelligence were positive or negative).

Participation coefficient was positively associated with individual intelligence scores in bilateral and medial prefrontal, bilateral parts of the IPL belonging to the TPJs, and in the posterior cingulate cortex (PCC). A negative association was only observed in one medially located paracentral region (Figure 5). The strongest effects that remained significant after correcting for the number of network nodes with FDR (200 cortical regions) were observed in a region of the left IPL belonging to the left TPJ (*rho* = 0.43, *p* = 0.033), a region of the right IPL belonging to the right TPJ (*rho* = 0.41, *p* = 0.039), in the left dorsolateral prefrontal cortex (*rho* = 0.48, *p* = 0.013), and in the right dorsolateral prefrontal cortex (*rho* = 0.42, *p* = 0.039; note that the latter two associations are only slightly visible in Figure 5 as located along the midline and are therefore highlighted by arrows). Again, similar results could be observed for varying functional connectivity thresholds (*Supplement* Figs. S5 and S6). Spatial correlations of intelligence-related aspects between both centrality measures are presented in the Supplement.

As frontal and parietal brain regions play a key role in established intelligence theories such as the PFIT^7^, we further explored whether the observed associations between intelligence differences and variations in participation coefficient are particularly localized to these regions compared to other regions. To this end, all Schaefer^48^ parcels (200 node Schaefer parcellation, see Methods) that anatomically overlap with frontal and parietal P-FIT regions were selected. Specifically, regions corresponding to the dorsolateral prefrontal cortex (BAs 6, 9, 10, 45, 46, 47), the inferior (BAs 39, 40) and superior (BA 7) parietal lobules, and the anterior cingulate cortex (BA 32), proposed by Jung & Haier^7^ were included. This mapping resulted in 42 of the 200 parcels. We then compared the mean correlation between participation coefficient and intelligence in these fronto-parietal parcels to the mean correlation of remaining parcels. Finally, a one-tailed permutation test (10,000 permutations) was used to assess statistical significance by randomly re-assigning parcel labels. The observed mean difference was significantly larger than expected under the null distribution (*p* = 0.002), providing empirical support for the hypothesis that frontal and parietal brain regions, as proposed by the P-FIT, show stronger associations compared to other cortical areas (see Supplementary Fig. S7).

### Intelligence is associated with multiscale entropy

In the EEG sample, MSE was significantly positively associated with intelligence at higher temporal scales (*p* < 0.05 by cluster-based permutation tests), while there was a trend towards negative associations at lower temporal scales at frontal electrodes (Figure 6). Significant clusters comprise brain-wide distributed electrodes except temporal electrodes. This pattern suggests that better performance in the intelligence test might link to more flexible long-range processes between a distributed network of brain regions.

**Figure 6.**
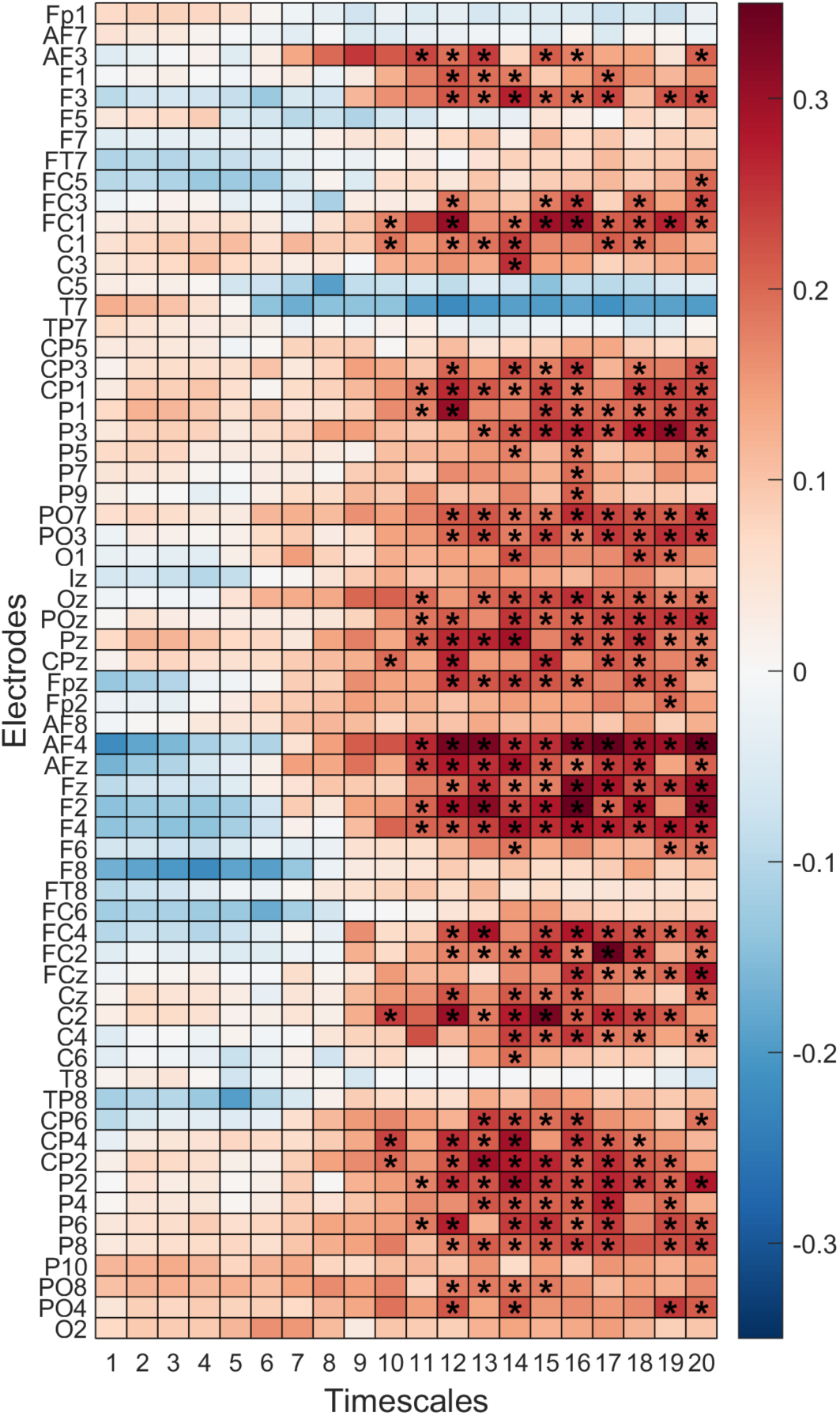
Spearman correlation between multiscale entropy (MSE) and intelligence during intelligence testing in Sample 2 (EEG). Each row corresponds to one EEG electrode, and each column to a timescale (1–20).

In Sample 2, each participant was presented with RPM items for a fixed duration of 30 seconds. However, the exact period during which participants actively worked on the items remained unknown. To address this, we examined associations between individual intelligence and MSE across different segments of the trial. Specifically, we analyzed the first, second, and third 10-second intervals following stimulus onset. As shown in Figure 7, the overall pattern of associations was consistent throughout the trial; however, the strongest associations emerged during the first 10 seconds, likely reflecting the period of highest cognitive engagement with the items.

**Figure 7.**
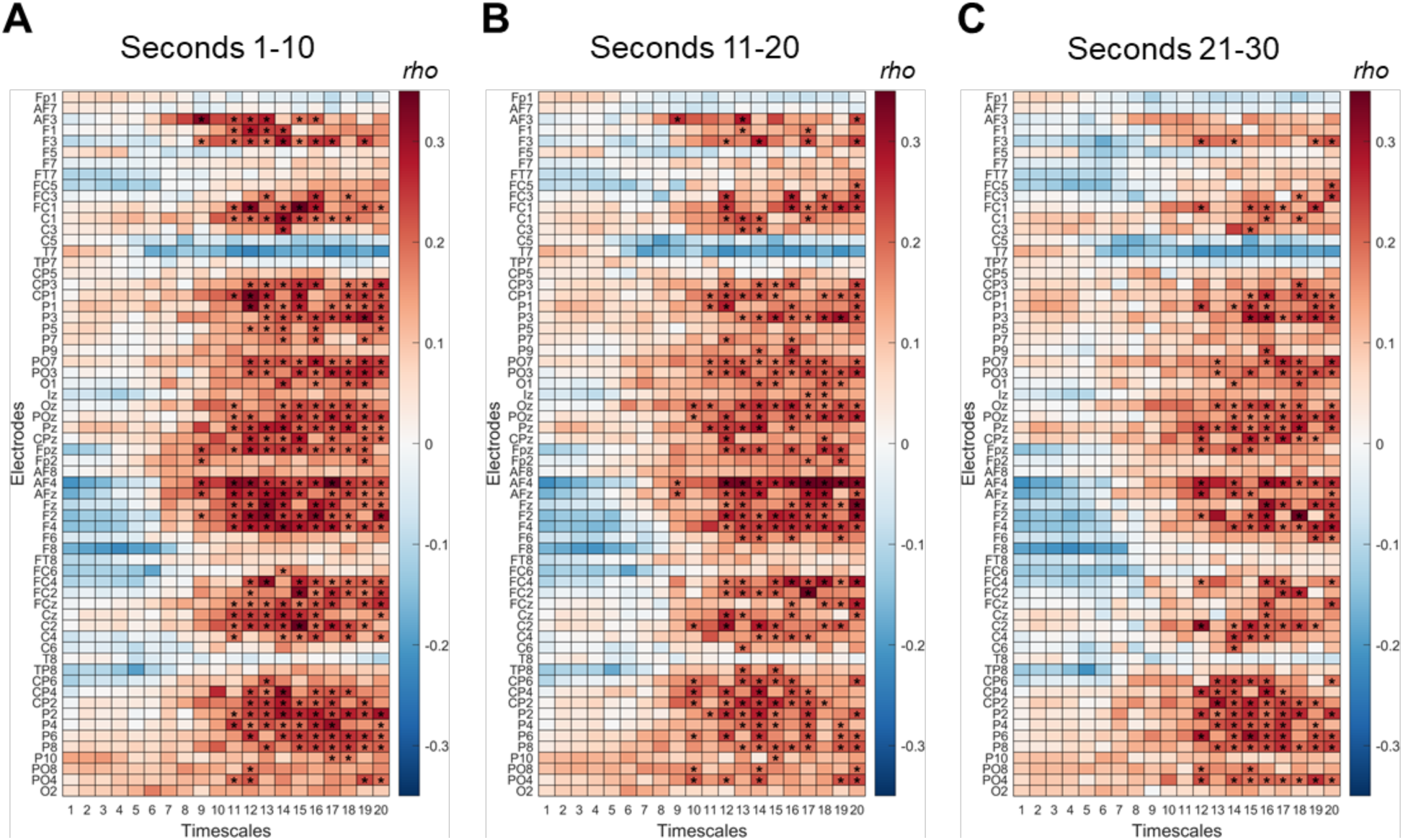
Spearman correlation between multiscale entropy (MSE) and intelligence during intelligence testing in Sample 2 (EEG) for the three consecutive periods (epochs) of the RPM trial. The first epoch is displayed in (A), the second epoch in (B), and the third (last) epoch in (C). Epochs lasted 10 seconds. Each row corresponds to one EEG electrode, and each column to a timescale (1– 20).

## Discussion

Our study builds on a large body of research relating individual differences in intelligence to characteristics of brain activity obtained during resting state or during rather simple cognitive tasks. Most of these studies employed either fMRI with limited insight into temporal dynamics or EEG providing restricted information about communication characteristics of brain regions involved in long-range processes. We extend this research by investigating intelligence-critical neural processes during behavior actually defining individual differences in intelligence - during performance of a well-established intelligence test. Specifically, we leveraged fMRI to explore how the overall connectedness of brain regions (degree) as well as their participation in intermodular communication (participation coefficient) relates to variations in intelligence. Additionally, we used EEG entropy analyses to shed light on the intelligence-critical neural complexity of processes at different timescales. While associations between intelligence and degree did not reach significance, we observed significant associations between individual intelligence scores and intermodular connectedness of specific frontal and parietal brain regions in the fMRI sample. These regions overlap with regions proposed by established intelligence theories and demonstrate the strongest reconfiguration in their connectivity pattern when switching from rest to task. In the EEG sample, intelligence was significantly positively associated with higher multiscale entropy (MSE) in brain-wide regions at coarser temporal scales. With this, our study allows for conclusions with respect to established localizationist theories like the P-FIT^6,7^, informs about newer network neuroscience proposals (NNT)^9^ and provides the first empirical test of the most recently introduced Multilayer Processing Theory (MLPT)^12^ that suggests higher intelligence as resulting from more flexible long-range processes at coarser time scales that more flexibly coordinate simpler short-range processes at finer timescales.

### Fronto-parietal long-range connectivity as driver of intelligence

While observing no significant associations between intelligence and the overall connectedness of brain regions (degree), relations between intelligence and the intermodular connectedness of brain regions were stronger and significant. This outcome suggests that not simply weaker or stronger connectivity of specific brain regions is critical to intelligence-related processing but highlights the diversity of connections with which those regions span across different functional brain systems. As all associations were positive, and higher diversity of connections results per definition from connecting different modules via long-range links, this observation provides support for the hypothesis of the MLPT that more flexible long-range processes relate to higher intelligence^12^.

The brain regions in which we observed significant associations comprise two regions of the bilateral dorsolateral prefrontal cortex located along the frontal midline and two parietal regions belonging to the bilateral TPJs. All four of these fronto-parietal regions are included in the brain network responsible for the default mode of human brain functioning (DMN)^49^ and overlap with regions, whose intrinsic intermodular connectedness has been related to intelligence previously^17^ as well as with brain regions proposed as relevant for intelligence by established theories such as the P-FIT^6,7^ and the multiple-demand theory^50^. Further supporting the P-FIT, we found that the effect was significantly more pronounced in frontal and parietal regions —those proposed by the theory—than in all other brain regions. As all significant associations were positive, our results suggest that fronto-parietal brain regions have more diverse connectivity patterns to other brain modules (beyond the fronto-parietal control network) in people with higher intelligence scores. Thus, while the P-FIT proposes efficient communication *within* the parieto-frontal control network as neural mechanism important to intelligence, the current study extends this proposal by suggesting that increased communication *between* the frontal-parietal network and other networks (*via* integrative long-range connections of prefrontal and parietal hubs to other modules) might be key for intelligence-related processing.

Interestingly, the same frontal and parietal regions were among those demonstrating the strongest reconfiguration in both centrality measures when switching from rest to task. In general, transition from resting state to task has been found to be accompanied by changes in functional organization from a more modular segregated brain state to a more integrated configuration^51^, i.e., communication between functional networks is increased, especially during cognitive tasks^52^. The DMN was proposed to play an important role within this process in acting as global integrator, facilitating activity in frontoparietal networks during the conscious processing of information^35,53–56^. Associations between the success in the intelligence test and the intermodular connectedness of prefrontal and parietal regions of the DMN might therefore be attributed to the role of those regions in increasing the communication between different brain networks via long-range processes and in transitioning into more integrated states, as required for solving complex cognitive tasks such as intelligence tests.

### Network flexibility as key mechanism

Leveraging EEG, we observed a significant positive association between better task performance in the RPM and higher multiscale entropy (MSE) at higher temporal scales — a finding that aligns with previous research on resting-state data^33^. Since brain signal complexity has been linked to the richness of information in the signal^26,27^ as well as to the flexibility of neural dynamics^28^, higher complexity of the neural signal might reflect more flexible changes in neural processes, which supports one tenet of the MLPT: namely, higher intelligence results from more flexible long-range processes at coarser timescales, coordinating short-range processes at finer timescales. According to the MLTP, these short-range processes (presumably reflected in finer timescales) are expected to be less complex in individuals with higher intelligence scores. Empirical support for such a negative association between complexity at finer timescales and intelligence has previously only been observed during rest^33,57^. Extending this, our data acquired during RPM performance, showed a trend-level negative association between intelligence and MSE at finer temporal scales at frontal electrodes. However, this effect did not reach statistical significance, providing only tentative support for this second prediction of the MLPT. Thus, future research should investigate this aspect in more detail, particularly considering the challenge of capturing short-range dynamics with limited spatial resolution. Critically, neural short-range processes are more susceptible to signal overlap and volume conduction as they are typically reflected in higher-frequency activity and confined to small neural assemblies. In contrast, long-range interactions—often associated with lower frequencies and broader spatial synchronization— can be more robustly captured by EEG. This aspect could have contributed to the clearer findings at coarser temporal scales.

Based on a) our finding of links between higher intelligence and higher complexity of long-range processes, b) our finding of higher intermodular connectedness of fronto-parietal regions, and c) previous research, we propose network flexibility as a key mechanism underlying successful intelligence test performance. In this framework, network flexibility refers to the ability of brain networks to adapt their connections efficiently to switch between cognitive strategies. Although our investigation of brain network characteristics was limited to static connectivity, we hypothesize that specific frontal and parietal brain regions revealed as critical for switches between rest and RPM might also be involved in the dynamic reconfiguration of brain activity states during performing the RPM, reflected in the association between intelligence and intermodular connectedness. This would align with the NNT, which proposes the flexibility to drive the brain network into different activity states as a neural foundation of general intelligence^9^ and assumes that switches in brain activity states are initiated and coordinated by frontoparietal regions through their connections to diverse brain networks^9,58^. More specifically, the NNT differentiates between easy and difficult-to-reach brain states in which the brain network transitions to meet specific cognitive demands^9^. Therein, while easy to reach states assumably support crystalized intelligence, difficult-to-reach states are needed for fluid reasoning. Accordingly, an individual’s ability to flexibly transition to difficult-to-reach states of brain activity could be considered crucial for succeeding at the Raven’s Matrices test, which primarily requires fluid thinking skills and was performed during the recording of brain activity in this study.

Interestingly, a relation between individual differences in intelligence and larger changes in functional connectivity to rare distant brain activity states during cognition, potentially reflecting such difficult-to-reach states, has recently been demonstrated in a study using dynamic network analysis^19^. Further, it has been suggested that efficient intermodular connectedness is required to initiate these changes into difficult-to-reach states and might be coordinated by specific brain regions^14^. Our fMRI results suggest that prefrontal and parietal brain regions, in particular the dorsolateral prefrontal near the frontal midline and the IPL part of the TPJ, might be critically involved in this coordination process, i.e., for transitioning the brain flexibly into required brain states including difficult-to-reach states.

### Intelligence-related processes across temporal and spatial scales

Our findings support the role of P-FIT regions in intelligence-related processes and highlight the importance of dynamic flexibility within long-range networks, as emphasized in theories such as the NNT. However, these theories predominantly focus on processes occurring at relatively coarse temporal scales, often overlooking the significance of faster neural dynamics and the interactions across different temporal scales. In contrast, EEG-based cross-frequency coupling studies have demonstrated links between intelligence and interactions across multiple temporal scales^45,59^, yet these approaches lack spatial precision.

Building on results of this study and the framework proposed by the MLTP, we suggest that studying individual differences in intelligence through the lens of neural processes occurring across both temporal and spatial scales—using methods such as magnetoencephalography (MEG)^60^—represents a promising direction for future research.

### Limitations and future directions

Our study has several limitations. First, our sample sizes were limited, critically diminishing the statistical power to detect small effects^61–63^. Also, we had no access to a sample in which we could test for independent replication. Therefore, results should be interpreted with caution and future studies with larger samples (and higher numbers of trials) are essentially required to study this subject with larger power. Second, fMRI and EEG data stem from different samples, preventing direct comparisons. Third, EEG analysis in general has poor spatial resolution and might therefore be limited in detecting fast-scale characteristics of small neural assemblies, especially required for testing aspects of the MLTP. Future studies should employ methods of high spatial and high temporal resolution in the same participants (e.g., using MEG). Fourth, the age range of both samples was restricted to young adults. Thus, a last suggestion to future study is the investigation of a broad age range: Combined with the analysis of structural brain networks, this may inform about hypotheses whether the higher diversity of intermodular connections observed in fronto-parietal brain regions may present an innate attribute or reflect previous experience; also, it could provide more insights into age-related changes of the complexity at different time scales^30^ and their link to the decrease of fluid intelligence with age^64,65^.

### Conclusion

Our study reveals that the diversity with which specific fronto-parietal brain regions communicate through long-range connections with different brain modules, as well as the complexity of long-range processes, are significantly associated with individuals’ performance on an established intelligence test. Both of these aspects —connection diversity and long-range complexity —show substantial reconfiguration when switching from rest to task and might be involved in coordinating the communication between brain networks in adaptation to increasing task demands, thus facilitating successful task performance.

Importantly, our findings provide the first empirical evidence for the key assumptions of the Multilayer Processing Theory (MLPT), which posits that higher intelligence emerges from more flexible global long-range processes operating at coarser time scales, coordinating simpler short-range processes within smaller neuronal assemblies at finer time scales. While the long-range properties central to the MLPT were robustly observed in both the fMRI and EEG samples, the theory’s claims regarding short-range processes warrant further research. Overall, this study demonstrates the value of investigating the neural underpinnings of individual differences in intelligence during intelligence test performance and should motivate researchers to explore different methodologies for assessing brain activity towards this goal.

## Methods

We used neuroscientific data sets from two independent laboratories: the first to assess characteristics of slow functional communication and to identify relevant brain regions; the second to consider different higher time scales containing information about the temporal dynamics of involved neural processes. The first sample comprises fMRI data from 85 participants of one resting-state run and three runs recorded during solving items from Raven’s Standard Progressive Matrices (RSPM) as well as items from Raven’s Advanced Progressive Matrices (RAPM)^66,67^, including data previously published by Vakhtin et al.^68^. These data were recorded under supervision of Rex Jung (University of New Mexico) at the Mind Research Network (MRN), Albuquerque, NM. The second data set comprises EEG data of 161 participants recorded during resting state and during Raven’s Advanced Progressive Matrices^67^ with 36 items. Data was provided by the research group of Adam Chuderski (Centre for Cognitive Science, Jagiellonian University in Krakow), including data published by Ociepka et al.^44^. Note that, due to the mix of RSPM and RAPM items in the first sample, for both samples, items are referred to collectively as RPM items in the subsequent text. Each RPM item presents a three-by-three matrix with eight elements of geometric shapes and one missing element. This missing element must be solved from the other elements that are linked to each other by logical rules.

### Sample 1

Sample 1 included 85 participants (34 females, age range: 18-29 years, *M*: 22.35; *SD*: 3.14) that were screened and excluded if they reported past major head injuries, psychiatric or neurological disorders, substance abuse, or consumption of any psychoactive medications. Informed consent was obtained from all participants. The study and consent form were approved by the UNM Institutional Review Board, and all procedures were implemented in accordance with the Declaration of Helsinki. After exclusions due to excessive head motion and short data length (see section Functional MRI Preprocessing and section Statistical Analysis), the final sample consisted of 67 participants (26 females; age range: 18-29 years, mean age: 22.91 years, age SD: 3.19 years; 56 right-handed, 9 left-handed, 2 with missing handedness data).

#### Stimuli and experimental procedure

Each RPM run included 10 items that were pseudo-randomly sampled from 30 items that were identical for each participant. Each item was presented for a maximum time of 15 seconds without showing the solution possibilities. Participants were instructed to push a button once they solved the item. After pushing the button or 15 seconds had passed a solution set appeared on the screen. Participants were then required to select the solution they believed to be correct by pressing one of four buttons, corresponding to the index and middle fingers of either hand. Only one response was permitted. Between each item, there was a period of 12 to 15 seconds during which a fixation cross was shown.

#### Functional MRI data acquisition

Functional MRI data were acquired using a 3T Siemens Trio scanner. Specifically, T2*-weighted functional images were obtained with a gradient-echo echo planar imaging sequence (echo time: 29ms, repetition time: 2s, 75° flip angle, 3.5 mm slice thickness, 30% distance factor, 240 mm field of view, voxel size: 3.8 mm × 3.8 mm × 3.5 mm). Resting-state runs lasted 5 minutes and 16 seconds, whereas RPM runs had a length of 5 minutes and 50 seconds.

#### Functional MRI preprocessing

Basic preprocessing of fMRI data was performed using FMRIPREP^69,70^ and is descibed in detail in the Supplementary Information. Briefly, preprocessing in FMRIPREP included intensity non-uniformity correction^71^, skull-stripping of each T1w volume as well as surface reconstruction^72^ and spatial normalization to the ICBM 152 Nonlinear Asymmetrical template^73^. Functional data were slice time^74^ and motion corrected^75^ which was followed by co-registration to the corresponding T1w using boundary-based registration^76^. Further preprocessing steps comprised high pass filtering with 0.008 Hz and a nuisance regression strategy according to Parkes et al.^77^ (pipeline no. 7) including 24 head motion regressors, and 10 components (aCompCor, five white matter and five cerebral fluid) from a principal component analysis (PCA) to putative nuisance signals^78^. For data recorded during the RPM, basis-set task regressors^79^ were applied simultaneously with the nuisance regressors to remove task-evoked neural activity, as task activation has been shown to produce systematic inflation of task functional connectivity estimates^79^. Finally, time series of nuisance-regressed blood oxygen-level dependent (BOLD) activity were extracted from 200 nodes covering the entire cortex^48^ (17 network parcellation^80^). Participants were excluded, if the mean framewise displacement (FD) exceeded 0.5 mm, any spikes above 5 mm were detected or if problem-solving times in the RPM runs (see section *Statistical analysis*) were shorter than resting-state runs.

#### Functional MRI-based functional brain connectivity

Functional connectivity between all 200 cortical nodes was estimated as Pearson correlation between z-standardized BOLD signal time series of a) the resting-state run, and b) the three concatenated RPM runs (including both correct and incorrect trials). Note that for the RPM runs, only those time-points were selected during which the RPM items were presented.

Those periods depended on the individual time a participant took to solve an item. For this reason, time points were filtered by taking the *n* highest points of the individual’s expected hemodynamic response, with *n* equals the duration of the presentation of the respective item. Further, the RPM data was shortened to fit the length of the resting-state trials (see section Statistical Analysis). Afterward, functional connectivity matrices were Fisher-z-transformed.

#### Graph-theoretical measures

Two graph-theoretical centrality measures, degree and participation coefficient, were calculated for each node using the MATLAB-based Brain Connectivity Toolbox (BCT)^81^ to capture a) the overall connectedness of a brain region within the network and b) this region’s involvement in connections linking different brain systems (diversity of intermodular connectedness). Both measures were determined on proportionally thresholded (50%) functional connectivity matrices. For testing whether the threshold critically affects results, all analyses were repeated with proportional thresholds of 40% and 60%.

The degree *k* of a given node *i* expresses how strongly this node is connected to all other nodes in the network and is calculated as sum of all weights (strengths) from adjacent connections, as follows:

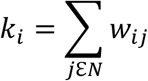

where *w_ij_* represents the functional connectivity strength between node *i* and node *j*, and *N* is the number of nodes in the network^81^.

The participation coefficient *y* of a node *i* assesses the diversity of its intermodular connections and is calculated based on modules within the functional network, as follows:

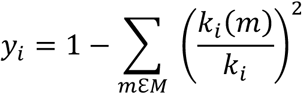

where *M* is the set of modules (clusters of nodes), *k_i_* is the node’s degree (see above), and *k_i_*(*m*) is the sum of strengths of connections between *i* and all nodes in module *m*^81^. For fMRI, nodes were assigned to the established fMRI-based partition of Yeo et al.^80^ including seven functional brain networks that served as network modules.

### Sample 2

Sample 2 included 161 right-handed participants (77 females, age range: 18-40 years, *M*: 23.60, *SD*: 4.28). All participants had normal or corrected-to-normal vision, and no history of neurological problems. The study was approved by the local ethics board, informed consent was obtained from each participant, and all procedures were implemented in accordance with the Declaration of Helsinki. Participants were excluded if demographic information (age or sex) was missing, as these were used as control variables. Additionally, participants with an insufficient number of usable trials due to artifact rejection were excluded (see section Statistical Analysis for details). This resulted in 131 participants (65 females, age range: 18-40 years, *M*: 23.63, *SD*: 4.31, all right-handed) remaining for analysis.

#### Stimuli and experimental procedure

The order of the RPM items was randomized for each participant. Each item was presented on a screen for 30 seconds without the solution possibilities. Afterward, the solution possibilities were shown, and participants had to select the solution they believed to be correct within a time window of 20 seconds. Only one response was permitted. Only EEG data recorded during the time in which RPM items but not solution possibilities were displayed was used for further analysis.

#### EEG data acquisition

Data were recorded in a sound-shielded cabin under supervision of an experimenter. First, 360 seconds of resting-state EEG were acquired followed by the RPM task. EEG was obtained using the Biosemi Active Two system with 64 Ag/AgCl scalp electrodes arranged in accordance with the 10-20 layout. Four further electrodes were placed above and below the left eye and in the external canthi of both eyes to capture vertical and horizontal electrooculograms (EOGs). Voltage values were referenced to zero (quantified relative to the driven right leg and common mode sense loop) and digitized at a sampling rate of 256 Hz.

#### EEG preprocessing

Preprocessing was done in MATLAB using EEGLAB^82^ (v2023.1). EEG data were re-referenced to the average of two mastoid electrodes. Next, a high-pass filter (1 Hz) and a notch filter (49.5-50.5 Hz) were applied, and ocular artifacts were corrected using the recursive least squares method^83^ as implemented in the EEGLAB plug-in ARR^84^. Following, resting-state data were segmented into 10 second and RPM data into 30 second epochs. Similar to the fMRI sample both correct and incorrect RPM trials were included. Channels exceeding amplitudes of ±160 μV were interpolated and epochs were removed if more than half the channels exceeded this threshold using the EEG-Plugin TBT^85^.

#### EEG multiscale entropy

Sample entropy^86^ measures the temporal irregularity of time series—here, the EEG signal— based on the probability that two sequences of consecutive data points of length *m*, which are similar within a specified tolerance, remain similar when extended by one additional data point (at length *m*+1). Multiscale entropy (MSE)^26,87^ extends this approach by analyzing entropy over multiple timescales. To calculate MSE, multiple coarse-grained time series are first constructed from the original EEG signal (voltage amplitudes in the time domain) by averaging data points within non-overlapping segments of length 𝜏, where 𝜏 represents the scale factor. Sample entropy is then computed separately for each coarse-grained signal. For each scale, template vectors of length *m* (or *m*+1), here *m* = 2, are extracted from the coarse-grained time series. Then, distances between all pairs of vectors of length *m* and between all pairs of vectors of length *m*+1 are computed separately using the Chebyshev distance metric. Two vectors are considered similar (matching) if their distance is less than or equal to a tolerance *r*, set to 0.2 times the standard deviation of the original signal. The quantities *B* and *A* represent the counts of matching vector pairs for lengths *m* and *m*+1, normalized by the total number of possible vector pairs (excluding self-matches). Sample entropy is then calculated as:

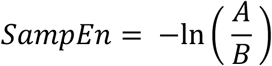

where a lower value of *SampEn* indicates more regularity, and a higher value indicates greater complexity or irregularity. Before computing MSE, EEG data was bandpassfiltered in the range of 1Hz – 40 Hz. For both RPM and resting-state trials, MSE was computed on 10 second epochs.

### Statistical analysis

To best assess RPM-related cognitive processing, individuals’ resting-state measures were subtracted from the RPM measures for both EEG and fMRI analyses. Intelligence was operationalized as sum score of the RPM, defined as the number of correctly solved items. Associations between brain measures and intelligence were determined as partial Spearman correlations controlled for age, sex (EEG and fMRI analyses), mean frame-wise displacement (fMRI analysis), and for the number of removed epochs due to artifacts and high occipital alpha (EEG analysis). Statistical significance was accepted for *p* < 0.05.

#### Sample 1

For most, but not all, participants, the RPM data comprised more time frames than the resting-state data. Therefore, to prevent statistical artifacts due to different data lengths, participants with fewer RPM time frames than resting-state frames were excluded. Further, the RPM data was shortened to fit the length of the resting-state trials (158 frames). Shortening the data was done by randomly selecting time frames to be excluded and was repeated 100 times. Following, centrality measures were averaged across these 100 iterations. For fMRI analysis *p*-values of node-wise correlations between the centrality measures and individual RPM scores were corrected via the false discovery rate (FDR, 200 nodes).

#### Sample 2

Because the number and length of trials varied across subjects, we standardized the data by equating the number of trials and epochs between conditions and subjects. Specifically, we included only participants who completed at least half of the RPM runs and had a minimum of 33 resting-state epochs. The minimum rest epoch count was chosen to allow for 11 RPM trials (each consisting of 3 epochs of 10 seconds), minimizing subject exclusion. For each participant, MSE values from RPM and resting-state data were averaged across selected trials and epochs. To ensure robustness, a permutation approach (100 iterations) was applied, randomly sampling RPM trials and resting-state epochs. The averaged MSE values across permutations were then used for subsequent correlation analyses with intelligence scores. Significance of correlations between MSE (64 electrodes x 20 timescales) and individual RPM scores were assessed by using a cluster-based permutation test^88^. In detail, clusters were defined as groups of spatially adjacent electrodes that each showed a significant positive or negative correlation (*p* < 0.05) at a given timescale. Adjacency was determined based on 3D Euclidean distances between standard 64-electrode locations, using a neighboring radius equal to half the mean inter-electrode distance. For each cluster, the test statistic was defined as the sum of correlation coefficients within the cluster. Then, intelligence scores were permuted across participants (1000 permutations). For each permutation, correlations were re-computed and the maximum positive and negative cluster-level statistics were extracted. Corrected *p*-values were then derived by comparing the observed cluster mean correlation to their respective permutation-based null distributions.

### Data and Code Availability

All data and analyses code has been made freely available. EEG data: https://osf.io/kv2sx (resting state), https://osf.io/htrsg (RPM), fMRI data and all analyses code: https://github.com/jonasAthiele/connectors_intelligence. Note, that fMRI data are shared in the preprocessed form, while raw data can be accessed from Rex Jung by reasonable request.

## Acknowledgements

This work was supported by the German Research Foundation (DFG, grant HI 2185/1-3 to K.H.), by the Heinrich-Böll Foundation (funds from the Federal Ministry of Education and Research, grant P145957 to J.A.T.), and by the Open Access Publication Fund of the University of Würzburg. Further, this research was supported in part by Lilly Endowment, Inc., through its support for the Indiana University Pervasive Technology Institute. This research was supported by a grant from the John Templeton Foundation (Grant # 22156) entitled “The Neuroscience of Scientific Creativity.” A.Ch. was supported by grant #2019/33/B/HS6/00321 from the National Science Centre of Poland.

## Author contributions

J.A.T.: Conceptualization, methodology, software, formal analysis, data curation, visualization, writing—original draft, funding acquisition. J.F.: Methodology, software, data curation, writing—review & editing. O.S.: Methodology, writing—review & editing. A.C.: Investigation, writing—review & editing. R.J.: Investigation, writing—review & editing. K.H.: Conceptualization, methodology, writing—original draft, supervision, project administration, funding acquisition.

## Notes

https://osf.io/kv2sx

https://osf.io/htrsg

https://github.com/jonasAthiele/connectors_intelligence

### Competing Interest Statement

The authors have declared no competing interest.

### Summary of Updates

The main changes include: 1. A revised EEG analysis approach by replacing static graph-theoretical centrality measures with dynamic multiscale entropy analyses. 2. Expanding the theoretical framing by integrating our findings with more recent theories of intelligence. 3. Conducting additional analyses to test regional specificity (e.g., frontal and parietal dominance), assess spatial correlations between centrality measures, and explore different time windows of the task trials. 4. Reporting only multiple comparison-corrected p-values.

