## Supplementary Information for "Decoding the Human Brain during Intelligence Testing"

### Supplementary Methods

#### Functional MRI preprocessing

Preprocessing of fMRI data was performed using FMRIPREP (Esteban et al., 2018, 2019). The following boilerplate text was automatically generated by FMRIPREP and ***users are advised to copy and paste this text into their manuscripts unchanged to avoid misspecifications***. It is released under the CC0 license:

Results included in this manuscript come from preprocessing performed using FMRIPREP version 20.0.7 (RRID:SCR_016216; Esteban et al., 2018, 2019), a Nipype (RRID:SCR_002502; Gorgolewski et al., 2011, 2018) based tool. Each T1w (T1-weighted) volume was corrected for INU (intensity non-uniformity) using *N4BiasFieldCorrection* v2.1.0 (Tustison et al., 2010) and skull-stripped using *antsBrainExtraction.sh* v2.1.0 (using the OASIS template). Brain surfaces were reconstructed using *recon-all* from FreeSurfer v6.0.1 (RRID:SCR_001847; Dale et al., 1999), and the brain mask estimated previously was refined with a custom variation of the method to reconcile ANTs-derived and FreeSurfer-derived segmentations of the cortical gray-matter of Mindboggle (RRID:SCR_002438; Klein et al., 2017). Spatial normalization to the ICBM 152 Nonlinear Asymmetrical template version 2009c (RRID:SCR_008796; Fonov et al., 2009) was performed through nonlinear registration with the *antsRegistration* tool of ANTs v2.1.0 (RRID:SCR_004757; Avants et al., 2008), using brain-extracted versions of both T1w volume and template. Brain tissue segmentation of cerebrospinal fluid (CSF), white-matter (WM) and gray-matter (GM) was performed on the brain-extracted T1w using *fast* (FSL v5.0.9, RRID:SCR_002823; Zhang et al., 2001). Functional data were slice time corrected using *3dTshift* from AFNI v16.2.07 (RRID:SCR_005927; Cox & Hyde, 1997) and motion corrected using *mcflirt* (FSL v5.0.9; Jenkinson et al., 2002). This was followed by co-registration to the corresponding T1w using boundary-based registration (Greve & Fischl, 2009) with six degrees of freedom, using *bbregister* (FreeSurfer v6.0.1). Motion correcting transformations, BOLD-to-T1w transformation and T1w-to-template (MNI) warp were concatenated and applied in a single step using *antsApplyTransforms* (ANTs v2.1.0) using Lanczos interpolation. Physiological noise regressors were extracted applying CompCor (Behzadi et al., 2007). Principal components were estimated for the two CompCor variants: temporal (tCompCor) and anatomical (aCompCor). A mask to exclude signal with cortical origin was obtained by eroding the brain mask, ensuring it only contained subcortical structures. Six tCompCor components were then calculated including only the top 5% variable voxels within that subcortical mask. For aCompCor, six components were calculated within the intersection of the subcortical mask and the union of CSF and WM masks calculated in T1w space, after their projection to the native space of each functional run. Frame-wise displacement (Power et al., 2014) was calculated for each functional run using the implementation of Nipype. Many internal operations of FMRIPREP use Nilearn (RRID:SCR_001362; Abraham et al., 2014), principally within the BOLD-processing workflow. For more details of the pipeline see https://fmriprep.readthedocs.io/en/20.0.7/workflows.html.

Further preprocessing steps comprised high pass filtering with 0.008 Hz and a nuisance regression strategy according to Parkes et al. (Parkes et al., 2018, pipeline no. 7) including 24 head motion regressors, and 10 components (aCompCor, five white matter and five cerebral fluid) from a principal component analysis (PCA) to putative nuisance signals (Muschelli et al., 2014). For data recorded during the RPM, basis-set task regressors (Cole et al., 2019) were applied simultaneously with the nuisance regressors to remove task-evoked neural activity, as task activation has been shown to produce systematic inflation of task functional connectivity estimates (Cole et al., 2019). Finally, time series of nuisance-regressed blood oxygen-level dependent (BOLD) activity were extracted from 200 nodes covering the entire cortex (Schaefer et al., 2018).

#### Supplementary Results

1. Correlation between Rest and RPM for participation coefficient (PC)

We observed a very high spatial correlation between the regional PC values during Rest and RPM ($r = 0.92$, $p < 0.001$), indicating that the majority of the diversity with which brain regions connect between modules stays consistent when switching from rest to task.

2. Correlation between Rest and RPM conditions for degree

Similarly, degree showed a strong spatial correlation between Rest and RPM ($r = 0.90$, $p < 0.001$).

Both findings are consistent with previous evidence showing that the brain's functional network architecture during task performance is largely shaped by an intrinsic network organization that is also present during rest, along with small task-general and task-specific reconfigurations (Cole et al., 2014).

3. Correlation between RPM-specific (RPM minus Rest) PC and degree

When comparing the spatial patterns of the difference maps (RPM minus Rest), we found a moderate correlation between PC and degree measures ($r = 0.68$, $p < 0.001$), suggesting partial overlap between both spatial centrality distributions. This is in line with previous evidence and was expected, as centrality measures are generally positively correlated (e.g., Hilger et al., 2017).

4. Spatial correlation between PC and degree for RPM associations

Focusing specifically on the associations between individual's PC/degree and individual's RPM scores, the spatial correlation between associations with PC and associations with degree was moderate ($r = 0.45$, $p < 0.001$), indicating that these connectivity metrics capture related but not identical intelligence-critical aspects of the functional connectivity architecture.

#### Supplementary Figures

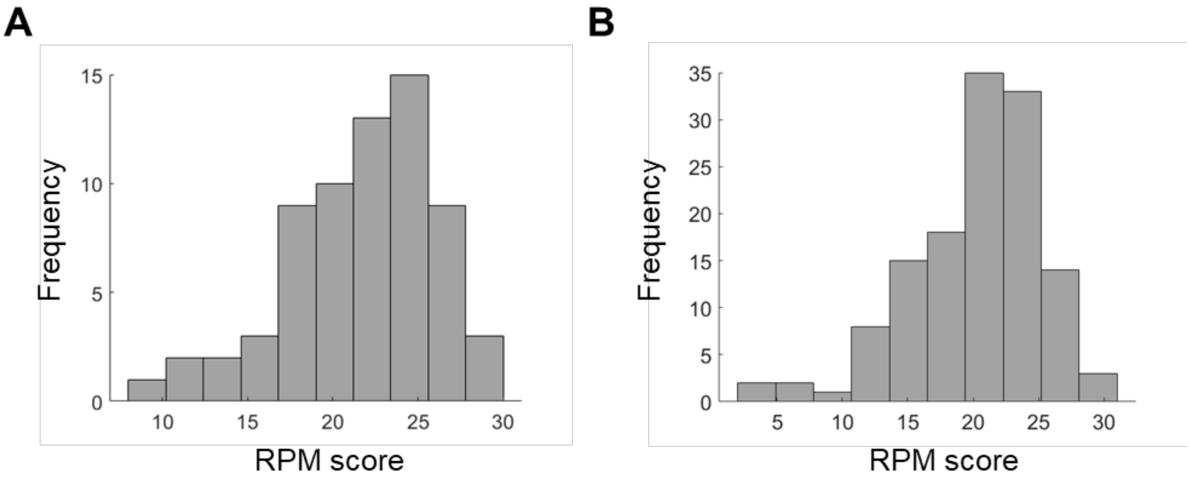


**Figure S1.** Distribution of intelligence scores (RPM sum scores). (A) RPM sum scores of the 67 participants of Sample 1. (B) RPM sum scores of the 131 participants of Sample 2. Note that, as only 30 items had to be solved in Sample 1, whereas the test in Sample 2 consisted of 36 items, we multiplied the original scores of Sample 1 by a factor of 36/30 to allow for better comparability. The scores illustrated in this Figure refer to the values after this adjustment.

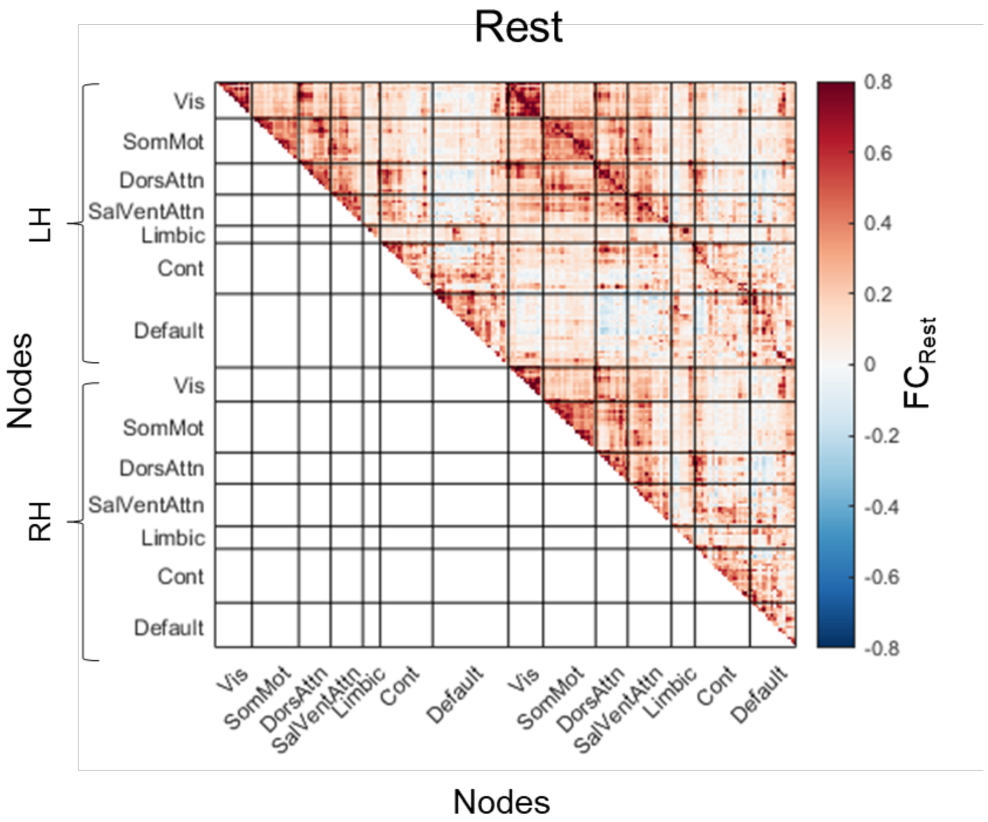


**Figure S2.** Functional connectivity derived from fMRI and averaged across all 67 participants of sample 1 during resting state.

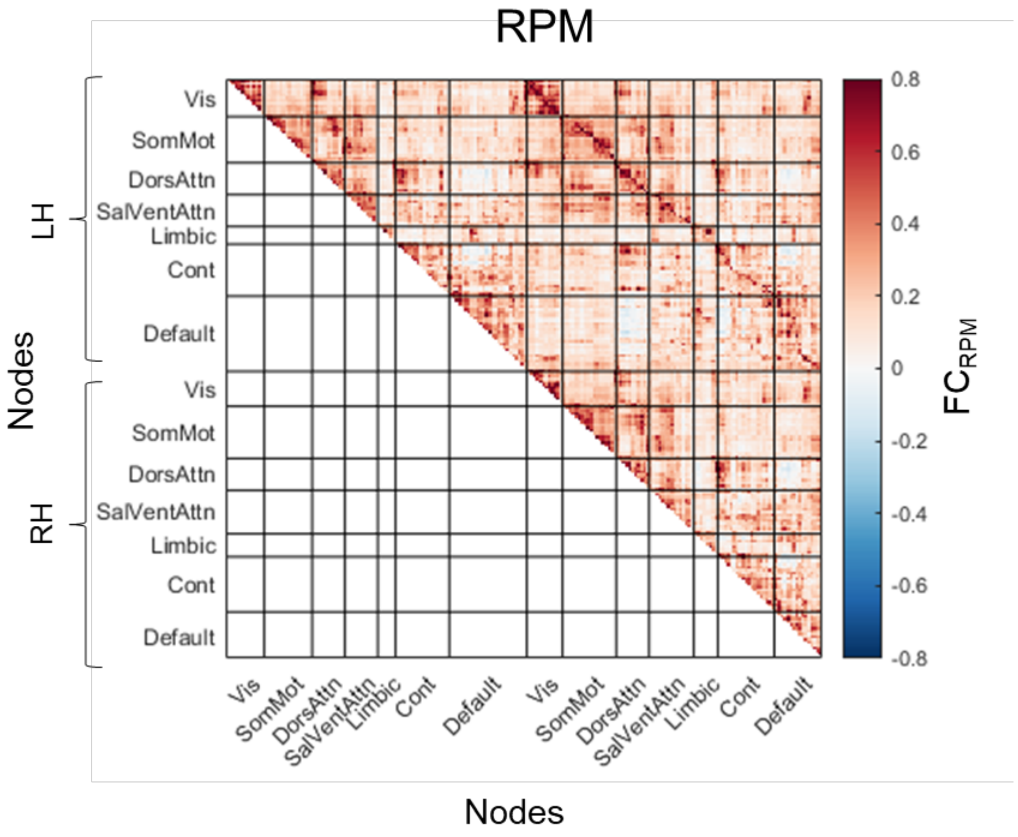


**Figure S3.** Functional connectivity derived from fMRI and averaged across all 67 participants of Sample 1 during RPM performance.

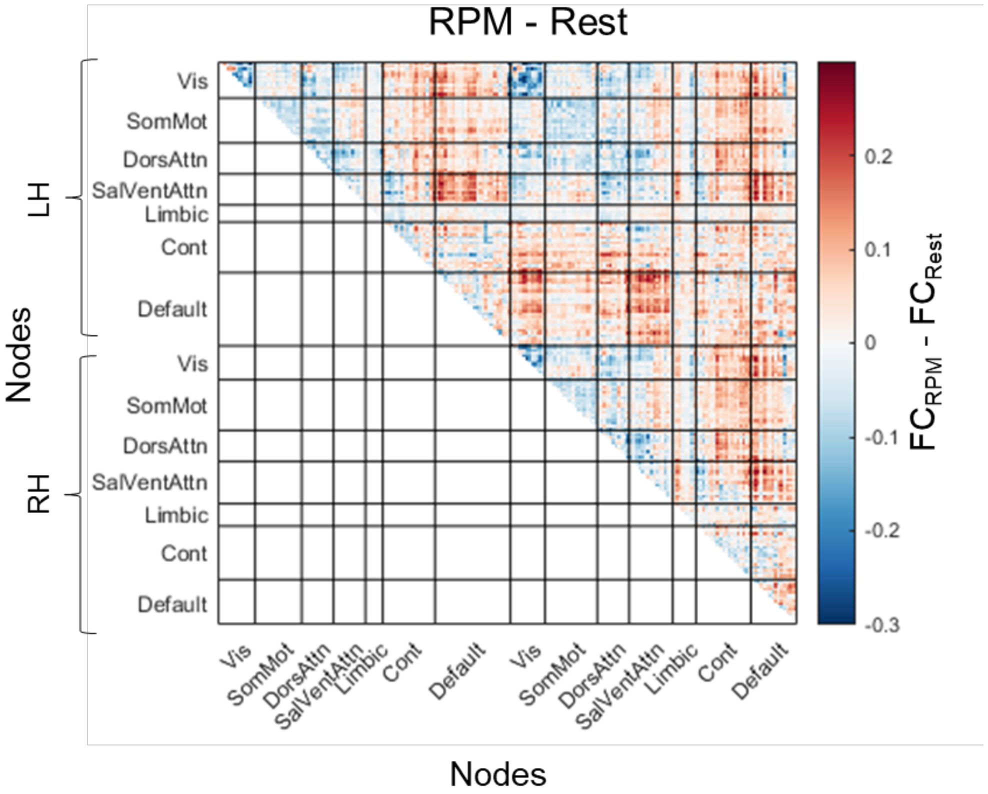


**Figure S4.** Difference in functional connectivity derived from fMRI between RPM performance and resting state averaged across all 67 participants of Sample 1.

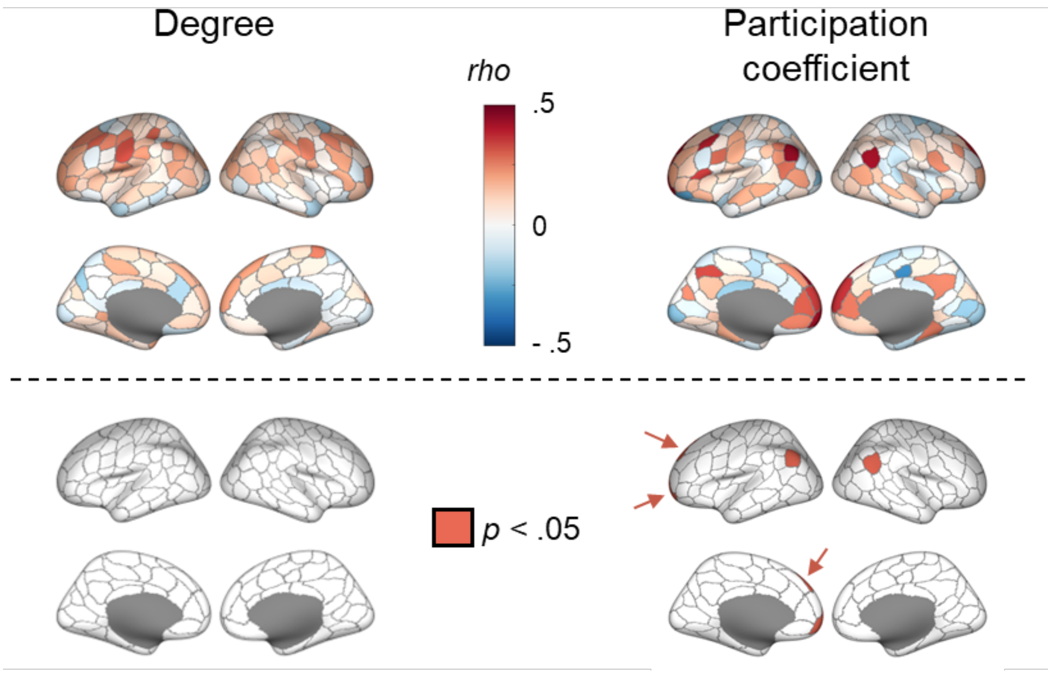


**Figure S5.** Spearman correlation between degree and intelligence during intelligence testing (left) as well as between participation coefficient and intelligence (right) in Sample 1 (fMRI) with a proportional threshold of 40% applied on the functional connectivity matrices. Upper panels show the Spearman correlation *rho* controlled for age, sex, and head motion. In lower panels, areas with $p$-values < 0.05, indicating significance (corrected for the number of network nodes with FDR, i.e., 200 brain regions) are colored in red (irrespective of whether associations with intelligence were positive or negative).

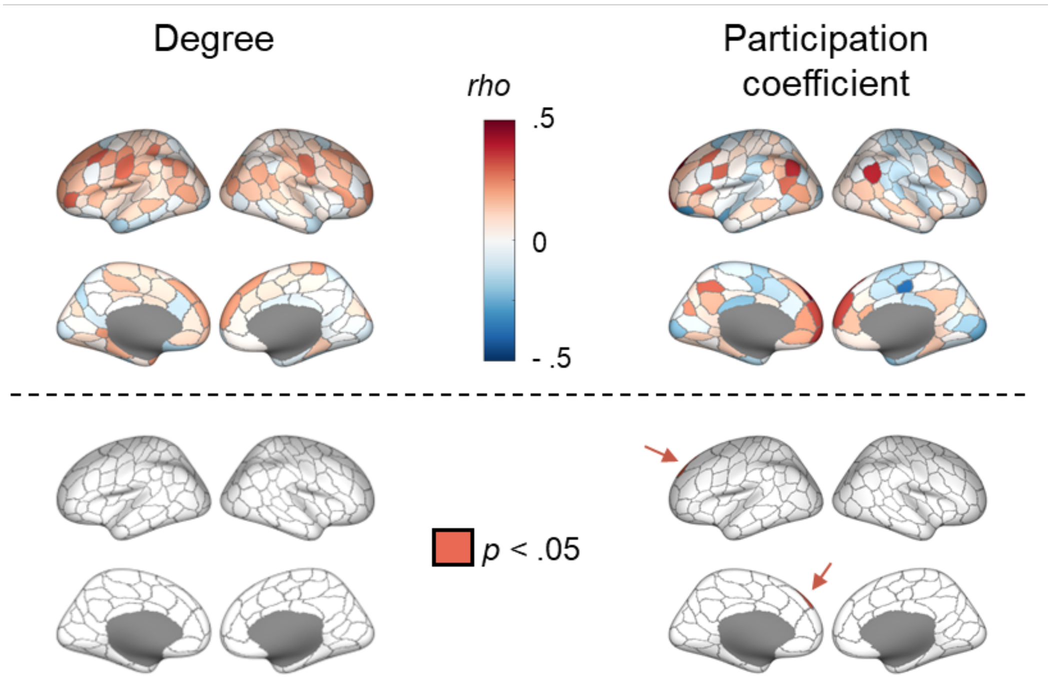


**Figure S6.** Spearman correlation between degree and intelligence during intelligence testing (left) as well as between participation coefficient and intelligence (right) in Sample 1 (fMRI) with a proportional threshold of 60% applied on the functional connectivity matrices. Upper panels show the Spearman correlation *rho* controlled for age, sex, and head motion. In lower panels, areas with *p*-values < 0.05, indicating significance (corrected for the number of network nodes with FDR, i.e., 200 brain regions) are colored in red (irrespective of whether associations with intelligence were positive or negative).

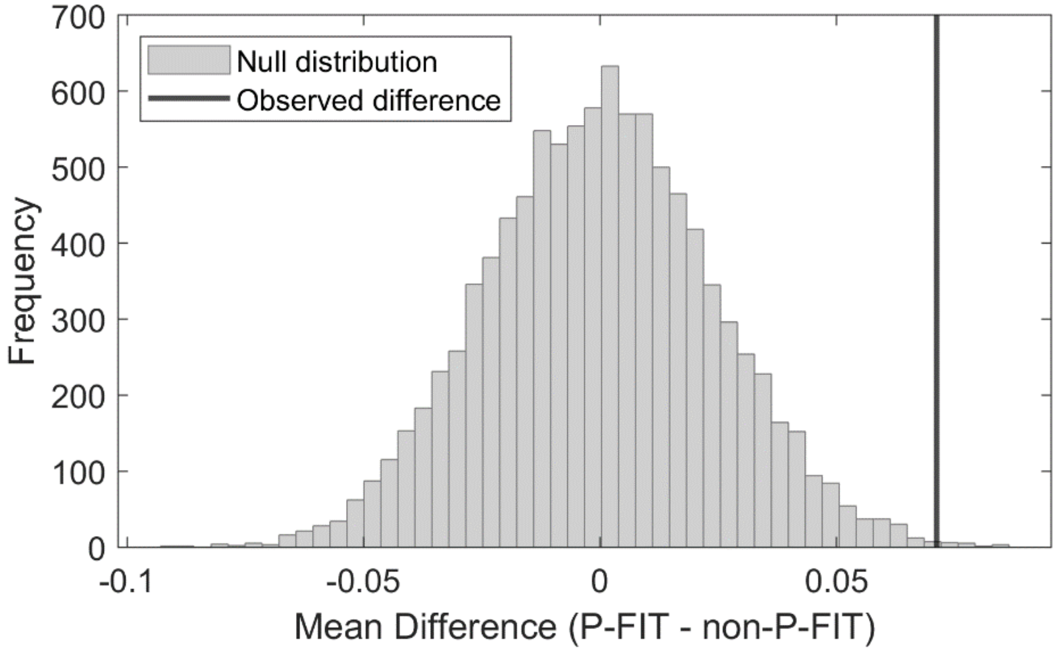


**Figure S7.** Permutation test null distribution of mean correlation differences between P-FIT and non-P-FIT regions. The histogram (grey bars) shows the distribution of mean differences generated by randomly shuffling parcel labels across 10,000 permutations. The thick black vertical line marks the observed mean difference (*rho* = 0.07), which lies significantly to the right of the null distribution, indicating stronger associations within P-FIT regions ($p$ = 0.002, one-tailed permutation test).
